# Perspectives on the design and performance of upper-limb wearable stimulation devices for stroke survivors with hemiplegia and spasticity

**DOI:** 10.1101/2020.08.20.260000

**Authors:** Caitlyn E. Seim, Brandon Ritter, Thad E. Starner, Kara Flavin, Maarten G. Lansberg, Allison M. Okamura

## Abstract

**Background:** Vibrotactile stimulation may improve limb function after stroke; however, current studies are limited by the stationary and clinic-based apparatus used to apply this stimulation. A wearable device could provide vibrotactile stimulation in a mobile form factor, enabling further study of this technique.

**Objective:** The aim of this work is twofold: (1) Design and validate a wearable device for stroke survivors that provides vibrotactile stimulation to the upper limb, and (2) Understand features that influence stroke survivors’ interaction with an upper-limb wearable device.

**Methods:** A vibrotactile glove was designed to apply stimulation to the stroke-affected hand. This device, the vibrotactile stimulation (VTS) Glove, features small electronics, provides hours of stimulation, and has the appearance of a fingerless fitness glove. We performed two studies. In Study 1, sixteen stroke survivors were given the glove and asked to wear it for three hours daily for eight weeks. Usage time, indicating adherence, was calculated by onboard sensors. The device was evaluated using a log of damages and feedback from participants. Self-reported behaviors during glove use were also collected. In Study 2, eight stroke survivors evaluated new device prototypes in a three-round iterative design study. Interviews were used to collect participant feedback as they donned and doffed the prototypes. Task completion time and correctness were measured. Thematic content analysis was used to define actionable design revisions between each round of evaluation and identify key perspectives on design principles.

**Results:** The VTS Glove is lightweight, wireless, and durable. Based on Study 1, adherence appears feasible. Each participant recorded over 140 hours wearing the VTS Glove. Participants reported wearing the glove during activities such as church, social events, dining out, and using the computer. However, 69% of participants struggled to extend or insert their fingers to don the device. Our analysis in Study 2 identified six other relevant perspectives from survivors regarding the wearable devices: hand supination is difficult, grip varies without awareness, devices may lead to sweat, separate finger attachments require dexterity, social comfort is perceived as unimportant, and the affected hand is infrequently used. New device designs with revised features (VTS Phalanx, VTS Armband, and VTS Palm) could be donned in an average time of 48 seconds (vs. 5.05 minutes to don the VTS Glove).

## Background and Motivation

There are approximately 6.6 million stroke survivors in the United States [1]. The American Heart Association projects that by 2030 there will be an additional 3.4 million Americans who have had a stroke (3.9% of the US population), because improved acute stroke care is leading to higher survival rates [1, 2]. Stroke often leads to physical disability, and upper-limb disability is a key factor in self-sufficiency. Following stroke, 35-55% of survivors have diminished tactile perception in their hand or arm [3, 4], and 50% have upper-limb motor disability [5, 6] (40-50% of whom also have spastic hypertonia [7, 8]). The most prevalent therapies available today are based on intensive limb use, such as constraint-induced movement therapy. However, for nearly 50% of stroke survivors this method is inaccessible due to low dexterity [5, 9] and adherence is a known challenge [10, 11, 12].

Stimulation-based methods of therapy may be more accessible than movement-based therapies to stroke survivors with low residual motor ability. Passively receiving stimulation presents less strain to survivors, including those who would not adhere to more taxing rehabilitation methods. Vibrotactile stimulation is used clinically to relieve muscle tone [13], and there is promising preliminary evidence that such stimulation may enable sensorimotor improvement and spasticity relief after central nervous system injuries like stroke and spinal cord injury [14, 15, 16, 17, 18, 19, 20, 21, 22, 23, 24, 25]. However, apparatus to apply this stimulation are not mobile and require assistance to operate. For example, two of the most prevalent methods are whole body vibration (WBV), which requires participants to stand on a scale-like platform, and repeated focal muscle vibration (rMV), which uses stationary machines with pins to provide stimulation. One device was used in laboratory work to deliver very low amplitude vibrotactile noise to the wrist, but provides only temporary feedback during task performance [26].

A wearable form factor would provide significant advantage to therapeutic vibrotactile stimulation. Since wearable devices can be closely coupled with the human body even while the wearer performs unrelated tasks, a wearable device could provide stimulation during daily life rather than during designated therapy time. A wearable form factor also allows longer durations and repetitions of vibrotactile stimulation, which have not yet been studied in clinical trials. Current laboratory studies typically apply this stimulation for periods of 5-30 minutes [16, 21, 24, 25, 27, 28, 29, 30]. More stimulation time may amplify results. Intensity (therapy time) is known to be associated with improved rehabilitation outcomes [31, 32, 33].

New wearable devices must be developed to support the study and implementation of vibrotactile stimulation therapy. The efficacy of such wearable devices depends both on the stimulation method and device use (adherence). Devices that are unused cannot result in therapeutic benefit. Thus, it is important to remove barriers to use before deploying new technologies. The Technology Acceptance Model discusses determinants of technology adoption [34, 35], the two cornerstones being ease of use and perceived usefulness. For wearable devices, these factors are influenced by breakage, dimensions and weight [36], perceived comfort [36, 37], and interaction time [35, 38, 39]. Well-designed devices allow clinical trials to focus on clinical efficacy, rather than adherence and logistical issues.

Wearable stimulation devices can be evaluated by addressing the following questions: Does the device deliver the required stimulation? Do patients wear the device as intended? The device must fit snugly to apply stimulation, be unobtrusive, and be durable for everyday wear. More importantly, the form of the device must be compatible with users who have upper limb disability. Wearable devices are like garments, and it is known that spastic hypertonia and hemiparesis can inhibit dressing even with the assistance of a caretaker [40, 41]. Accommodations that let affected individuals put on (don) and remove (doff) the device themselves enable participation in therapy for those without care-takers and independence for all involved. Donning and doffing are the primary interactions users have with a passive stimulation device; thus these interactions are a driving factor in the perceived ease of use for the device. Understanding how to reduce the time, difficulty, and confusion that may occur during these tasks allows these barriers to be addressed in advance by system designers.

Here we present the design of a lightweight, wireless, wearable device (called the VibroTactile Stimulation (VTS) Glove) that was designed to provide mobile vibrotactile stimulation to the disabled hand in chronic stroke. The VTS Glove was deployed in an eight-week study to understand if this method of therapy is feasible and assess device performance. Because participants experienced challenges donning and doffing the device, we performed a second study to find design accommodations and better understand factors that influence stroke survivors’ experience with an upper-limb wearable device.

### Preliminary Device Design: the VTS Glove

The first device design is a computerized, fingerless glove that delivers vibrotactile stimulation to the fingers of the affected hand.

#### Form Factor

Related work used vibrotactile gloves that synchronize with music to provide haptic training on how to play piano [42, 43]. In one study, these “MusicTouch” gloves were used to teach piano to patients with diminished hand dexterity from partial spinal cord injury [44]. The MusicTouch glove had some features suitable for able-bodied users and partial spinal cord injury survivors, namely: the glove was fingerless for fit, and palmless for sanitation and to allow wearing while operating a manual wheelchair [45]. The fingerless glove design worked well for this population – who had mild motor dysfunction (with no spasticity and hypertonia) from partial spinal cord injury. Therefore, a fingerless glove design was used when creating a preliminary VTS device for stroke survivors (Figure 1a). Flexfit Lifting gloves (Harbinger, Implus LLC.) were adapted to provide the base fabric for the prototype. The device is also lightweight, and can be used wirelessly. The appearance resembles exercise gloves, a recognizable form factor that may reduce confusion regarding how to wear the device correctly.

**Figure 1.**
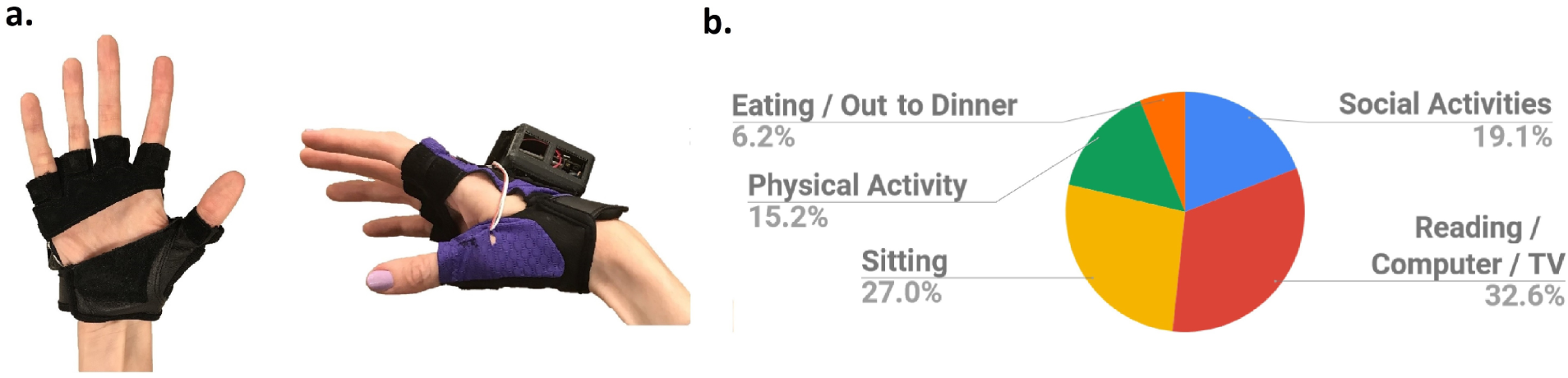
**(a**.) The VTS Glove, which provides wireless and wearable vibrotactile stimulation. **(b**.) Breakdown of activities performed while wearing the stimulation device, reported by sixteen stroke survivors in Study 1.

#### Electronics

Small vibration motors attach inside each finger sleeve of the glove and provide stimulation to the dorsal proximal phalanx. Reinforced wires run from these actuators to a circuitry box. Stimulation settings vary by individual study aims. In our preliminary design, the target was the cutaneous mechanoreceptors, specifically the Pacinian corpuscles, which respond to both direct vibration and vibration transmitted through the body at a frequency range of at least 10-400 Hz (preferentially responding at about 250 Hz) [46]. Coin-shaped, eccentric rotating mass (ERM) motors (Model #310-113, Precision Microdrives) were chosen for Study 1. These motors were driven with 3.3V current to produce an amplitude of 1.5 g and 210 Hz vibration frequency (measured in a laboratory setting for validation at 1.3 g and 175 Hz when attached to the glove). The glove stimulates each finger by activating the attached motor. The timing and pattern of stimuli across different motors is controlled by a microcontroller.

The circuit board contains a microcontroller, power regulator, USB port, and lithium polymer battery charging circuit, microSD card slot, motion-sensing chip, and motor-driving circuit. The microcontroller (Feather M0 Adalogger, Adafruit Industries) runs the device via preprogrammed software. The development board includes an onboard power regulator chip that ensures constant voltage of 3.3V. Thus, even when the battery ran low, stimuli for the patient remain uniform. A 50 mAh lithium polymer battery and charging circuit allow users to plug the glove into the wall to charge, then use the glove wirelessly for up to four hours. A triple axis gyroscope (L3GD20H, STMicro) detects vibration and movement, then logs this data to the native SD card. Data capture on the device is limited to timestamps and movement data. Security is handled by anonymization, and networking (such as Wifi or Bluetooth) is not part of this design. The motor-driving circuit uses Darlington transistor array chips (ULN2003, Texas Instruments Inc.) to draw current from the battery. This circuit is suitable for ERM-type motors. The electronics interface is simply a tactile switch that turns the stimulation on or off.

A 3D-printed box houses the glove’s control circuitry and secures to the back of the glove using Velcro (Figure 1a). This box rests on the rigid metacarpal bones of the hand, so as not to impede bending of the knuckles or wrist. The average length for the smallest adult metacarpal bone (pinky) is 2.12 inches [47]. The hardware components within the box have a footprint of 2.5 in. × 2.0 in.

## Methods: Study 1

VTS Gloves were given to stroke survivors to use at home for eight weeks. The aim of this study was to assess if adherence to this method of therapy is feasible and to gather data on device performance. Preliminary clinical data was also collected in this study and is presented in detail in a separate manuscript [48]. This trial provides essential new data on device performance in a home setting.

### Participants

Participants were sixteen chronic stroke survivors (ages 28-68, 1-13 years post stroke, 11 male/5 female) with diminished tactile perception and range of motion their hand. The study was conducted between August 2017 and February 2019. All participants provided written consent, and the protocol was approved by Georgia Institute of Technology’s Institutional Review Board.

### Study Design

Participants were each given a glove and asked to wear it switched on for three hours daily for eight weeks while awake. The hours did not need to be consecutive, and the device could be worn throughout the participant’s daily life. Participants were told that if the device was used less than 18 hours per week for two weeks, the participant would be counted as a drop-out. Although adherence under real-world conditions cannot be forecasted, this structure provides data on the potential for adherence to the device and stimulation routine. Participants used the glove for eight weeks and met with a study proctor weekly.

Half of the participants were randomly assigned using a lottery system to receive vibrotactile stimulation from the device, while the other half had vibration disabled. These conditions provide data on the impact of the stimulation. Instruction matched for both groups and device appearance was the same. Participants were asked only to use the device and were blinded to the different conditions.

### Measures

We hypothesized that stroke survivors would adhere to wearing the device daily and would find the VTS Glove tolerable with and without stimulation enabled. We also gathered participant feedback, to provide data on perceived ease of use and technology acceptance.

#### Usage Time Log

The glove automatically recorded participation time each week as a measure of adherence. Participation time was measured as the amount of time when the device was switched on. The total time when the device was switched on *and moving* was also recorded when the change in gyroscope values surpassed a predetermined threshold, binned into 10 second intervals. These data were logged to the microSD card and examined during weekly study visits. Proctors recorded if each total was or was not between 18-24 hours for the week.

#### Hardware Log

Areas of damage were logged by proctors as a measure of durability. In the event of damage, the device was repaired during a study visit or the participant was given a duplicate device.

#### Worksheet

Participants were given a worksheet each week to record the activities they performed while wearing the VTS Glove. The worksheet contained seven sections for notes, each labeled with a weekday. The second half of this worksheet included daily sections to record observations and comments. Participants were asked to write notes about device design, challenges, breakage, or anecdotes. Participants were also asked to record their self-reported daily wearing time on the worksheet.

#### Time to Don

During this study it was observed that participants struggled to get the glove on and off their affected hand. Thus, between their fifth and sixth week of participation, three participants were timed during the donning process.

## Results

### Adherence

All participants recorded at least the minimum of 18 hours of usage time each week, and none surpassed 24 total hours in a single week. No difference was found between groups. Total time when the device was on matched the total time when the device was on *and moving*, suggesting that the glove was not set aside while switched on.

### Hardware

The VTS Glove circuitry remained intact and successfully produced mobile stimulation for the duration of the study. Damage recorded during the study was limited to breaks in solder joints on the peripheral wires leading to the vibration motors. This likely occurred when wires were pulled during the don and doff process. Participants re-charged the devices daily using a wall adapter similar to other consumer electronics. Participants did not report confusion about how to wear and charge the device. The glove secured the vibration motors against the skin, though fit depended on custom sizes.

### Activities

Responses on the worksheet were analyzed by two independent raters using thematic content analysis. Then, themes were defined using affinity diagramming. Activities during VTS Glove use were associated with five themes which are reported in Figure 1b. Together, the themes of sitting and consuming media accounted for 59.6% of the reported activities. Some participants described wearing the glove to activities such as movies, church, brunch, family gatherings, and outdoor picnics.

### Challenges with don and doff

Eleven participants reported difficulty with the don and doff process on their worksheets. The VTS Glove could be worn with fingers in a tight grip – there were no reports of discomfort, sores or pressure ulcers. In addition, the glove can remain securely attached at the wrist if the fingers are open. However, the don and doff process was a challenge. Participants with spasticity or weakness had difficulty placing each finger in one of the glove sleeves, even with help from a caretaker. They **could not extend or insert their fingers**. Instead, their able hand was used to manipulate the affected fingers for extension and insertion. This prevented use of the able hand to simultaneously don and secure the device. The three randomly selected participants recorded an average time of 303 seconds to don the glove after participating in the study for at least four weeks.

## Methods: Study 2

We performed a second study to address the challenges with don and doff found in Study 1. The aim of this study was to create improved devices, and to identify the factors that influence stroke survivors’ experience using an upper-limb wearable device.

### Study Design

An iterative design-and-test methodology was used to evaluate three new prototypes. This structure enables multiple rounds of revision and introspection that cannot be achieved in a single-round study. (This is in contrast to a evaluation study using ranking measures such as the NASA TLX [49]). We hypothesized that, with revision, the new device prototypes could be donned in less time than the VTS Glove.

There were three rounds in this study. Two or three stroke survivors participated in each round. Between each round, the prototypes were revised. Each study visit was one hour in length. During their visit, each participant was asked to evaluate the prototypes one at a time in a randomized order. All procedures were repeated for each prototype. Each visit was recorded using a video camera and later transcribed verbatim. Study investigators had no relationship to the participants. Participants were assigned an alphanumeric code to anonymize study data.

### Participants

Participants were eight individuals with chronic stroke (ages 48-89, 6 months to 4 years post stroke, 5 male/3 female). Stroke survivors with diminished hand function were recruited through a local clinic according to the following inclusion criteria: history of stroke, no individual finger control on their affected hand, and limited range of motion in the fingers (<90° of voluntary extension at MCP joints). Individuals with aphasia were excluded because they could not provide verbal feedback. The setting was a university research lab, and the study was conducted in August 2019. All participants provided written consent, and the protocol was approved by the Stanford University Institutional Review Board.

### Design Goals

Design goals were identified for the new prototypes. Goals G1 and G2 were based on the results of Study G3-G6 are general features of wearable device design relevant to both able-bodied users and users with motor impairments.

#### G1: Fast to Don/Doff with Hand Spasticity or Weakness

Participants in Study 1 struggled to don the glove onto their affected hand. Reducing the interaction time is known to improve user experience and increase adoption of new technologies [34, 50]. We aim to reduce the time it takes to put on and remove the device, even when individuals have clenched hands due to spasticity and hypertonia, or limp fingers due to hemiparesis.

#### G2: Reliance on One Hand

Stroke affects primarily one side of the body, leading to diagnoses such as hemiparesis or hemiplegia [51]. When function is diminished on the affected side, the able hand is responsible for interacting with and adjusting equipment such as a wearable device. Not all users have a caretaker to assist them. A design that allows stroke survivors to operate the device independently (using only one hand) is more equitable and efficient. Secondly, the able hand is supported by the affected hand during some activities of daily living (ADLs). A wearable device should not significantly obstruct these activities of daily life.

#### G3: Easy to Use and Understand

Device design can encourage correct use. Correct use includes: how tight the device is worn and if the parts of the body are inserted correctly. Reducing confusion can also improve perceived ease of use, which is associated with technology acceptance [35].

#### G4: Physically and Socially Comfortable

Comfort of a wearable computing device includes physical and social components, because the device may be worn throughout daily life [52]. If users find a device uncomfortable, they are less likely to wear it [36].

#### G5: Fit

A one-size-fits-all, adjustable device allows more rapid production than custom devices or multiple sizes. A good fit also enables sensors and actuators to make reliable contact with the skin.

#### G6: Durability

For our application of wearable vibrotactile stimulation, devices may be worn for extended periods of time during the user’s daily activities. Repeated donning/doffing will result in strain on the user and wear on the device, so designs must be robust. If devices are used in rural or underserved areas, they must remain functional or be easily repaired.

### New VTS Device Designs

Based on the design requirements, three candidate designs were created for use in Study 2. One design, the VTS Phalanx, is a revised form of the VTS Glove.

Two additional designs, the VTS Armband and the VTS Palm, were also created. These additional designs address the opportunity to apply vibrotactile stimulation to different regions of the upper limb. Some research suggests a therapeutic benefit from stimulation at the muscle belly [21, 16], while others apply cutaneous stimulation [53]. The VTS Armband wraps over musculotendinous regions of the forearm. The VTS Palm device attaches to tissue that is dense in cutaneous sensory receptors (the glabrous skin of the volar hand) [54]. None can fully isolate a single type of tissue while maintaining a wearable form factor, but each of these designs form the basis for further research on stimulation at different locations.

Each design aims to apply four or more coin-shaped actuators to the body at the designated location (though no stimulation was applied during Study 2). All were designed to dissuade insertion or extension of the fingers, in view of Study 1 results. Each device design was fabricated into a non-stimulating prototype to allow participants to interact with design features. Prototypes also included an acrylic box at the location where electronics can be mounted.

#### VTS Phalanx

The VTS Phalanx is a revised form of the VTS Glove. Like the VTS Glove, this device (Figure 2a) is intended to provide vibrotactile simulation to the dorsal proximal phalanx. Fabric attaches with Velcro around the distal forearm and extends to the dorsal phalanges, leaving the palm exposed. This design maintains a similar shape to the VTS Glove; however, the glove’s finger sleeves are replaced with two straps that wrap over the fingers. These straps can be stretched over flexed fingers, so the design can be donned without finger extension. A rigid arch rests on each dorsal phalanx to keep the fingers in place without inserting the fingers individually. These arches are mounted on a foam bar that flexes to accommodate a variety of hand sizes.

**Figure 2.**
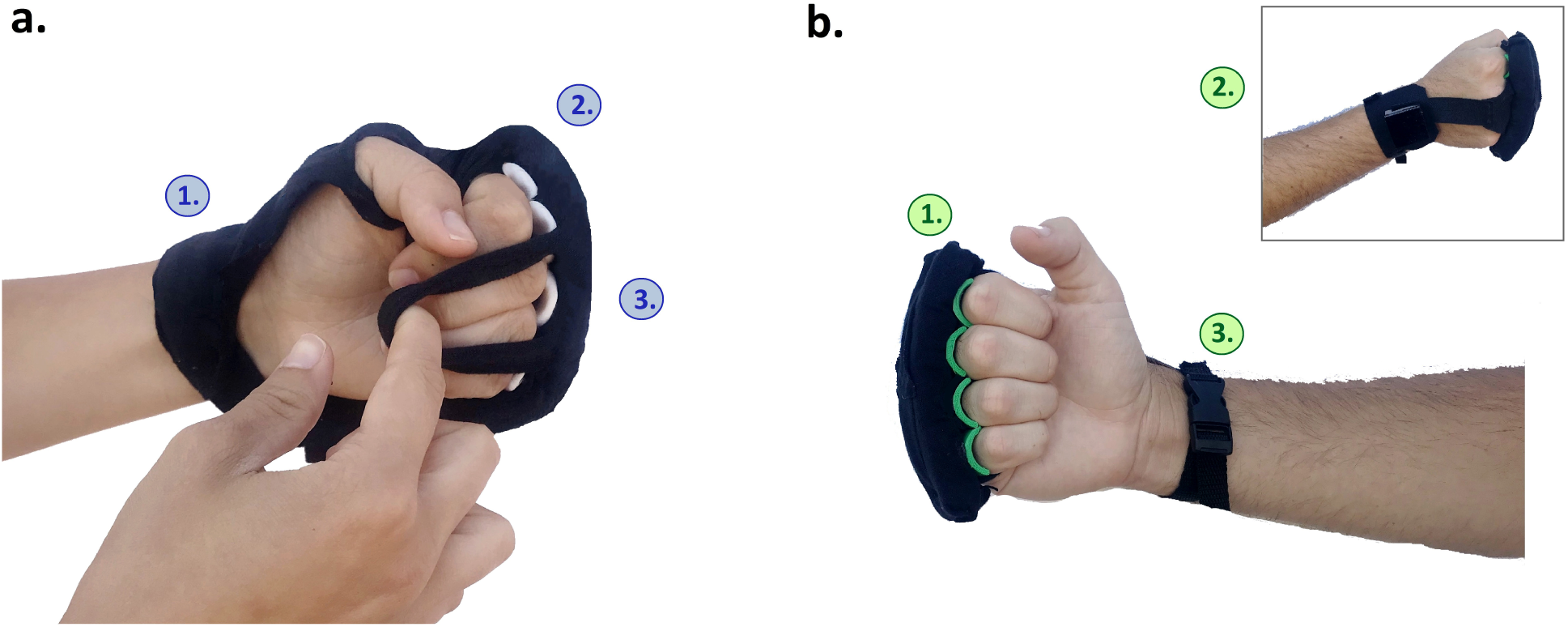
VTS Phalanx version V1 and version V4. **(a**.**) 1**. A form factor resembling fingerless gloves with the palm exposed. **2**. Rigid arches create a place for each finger without inserting each finger individually. **3**. The two straps are capable of wrapping over flexed fingers without requiring finger extension. **(b**.**) 1**. The single strap over the fingers can be donned with the anterior arm facing away (no supination). **2**. A T-strap design reduces fabric coverage while providing a path for wires to run between circuitry at the wrist and actuators at the fingers. **3**. A buckle closure enables strap adjustment using one hand.

#### VTS Armband

The Armband (Figure 3a) wraps around the forearm and allows stimulation to be applied to the muscle belly of the extrinsic hand muscles. The 3 inch-wide band is 15 inches long for fit around different size arms. The fabric band is attached using Velcro, and electronics can be mounted on or inside the band.

**Figure 3.**
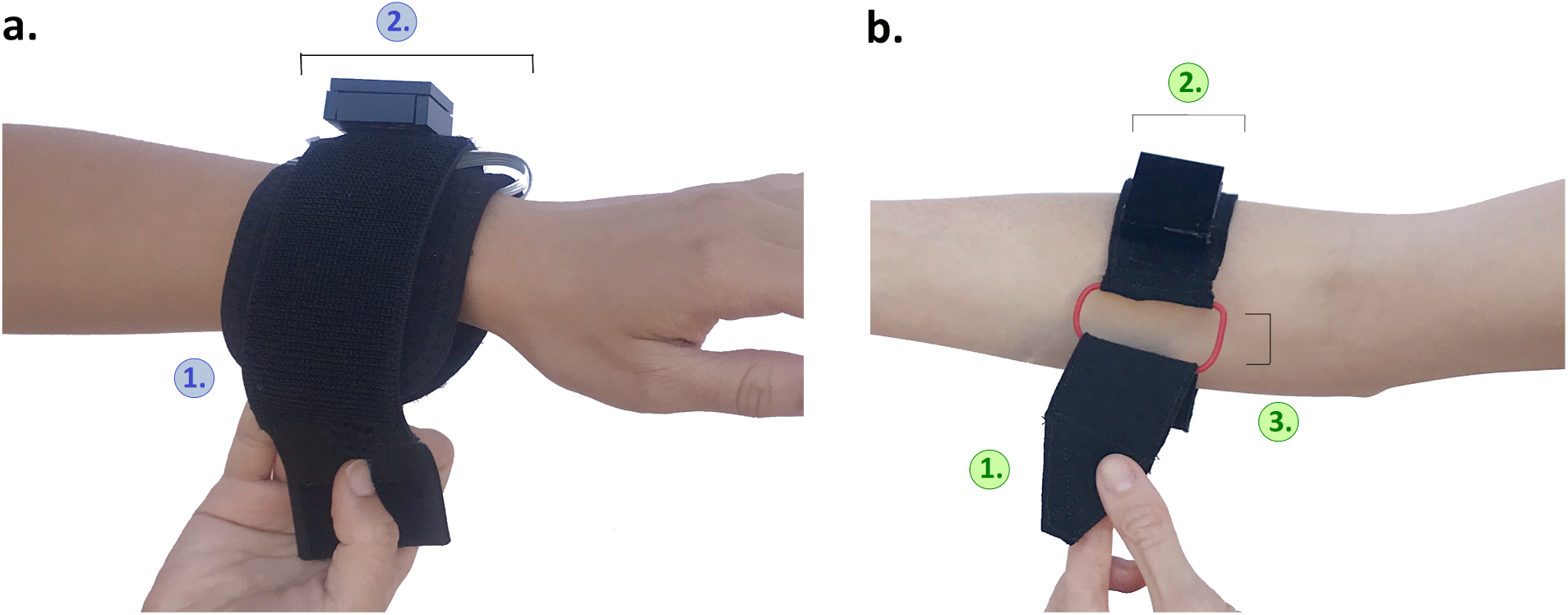
VTS Armband version V1 and version V4. **(a**.**) 1**. The rectangular armband wraps around the arm and attaches via Velcro. 2. The band has a width of 3 inches and circuitry can be mounted on or in the armband. **(b**.**) 1**. The rigid and tapered end of the armband is designed to slip through the 0.75 inch buckle frame. **2**. The armband is 2 inches wide. **3**. A cinch buckle enables users to secure the device using one hand.

#### VTS Palm

This design (Figure 4a) enables stimulation at the volar hand (palm) and is intended to be grippable. This prototype has a graspable form factor. The body of the device is a foam rod (4 inches). The foam rod is reinforced inside with a semi-rigid tube that provides structural support.

**Figure 4.**
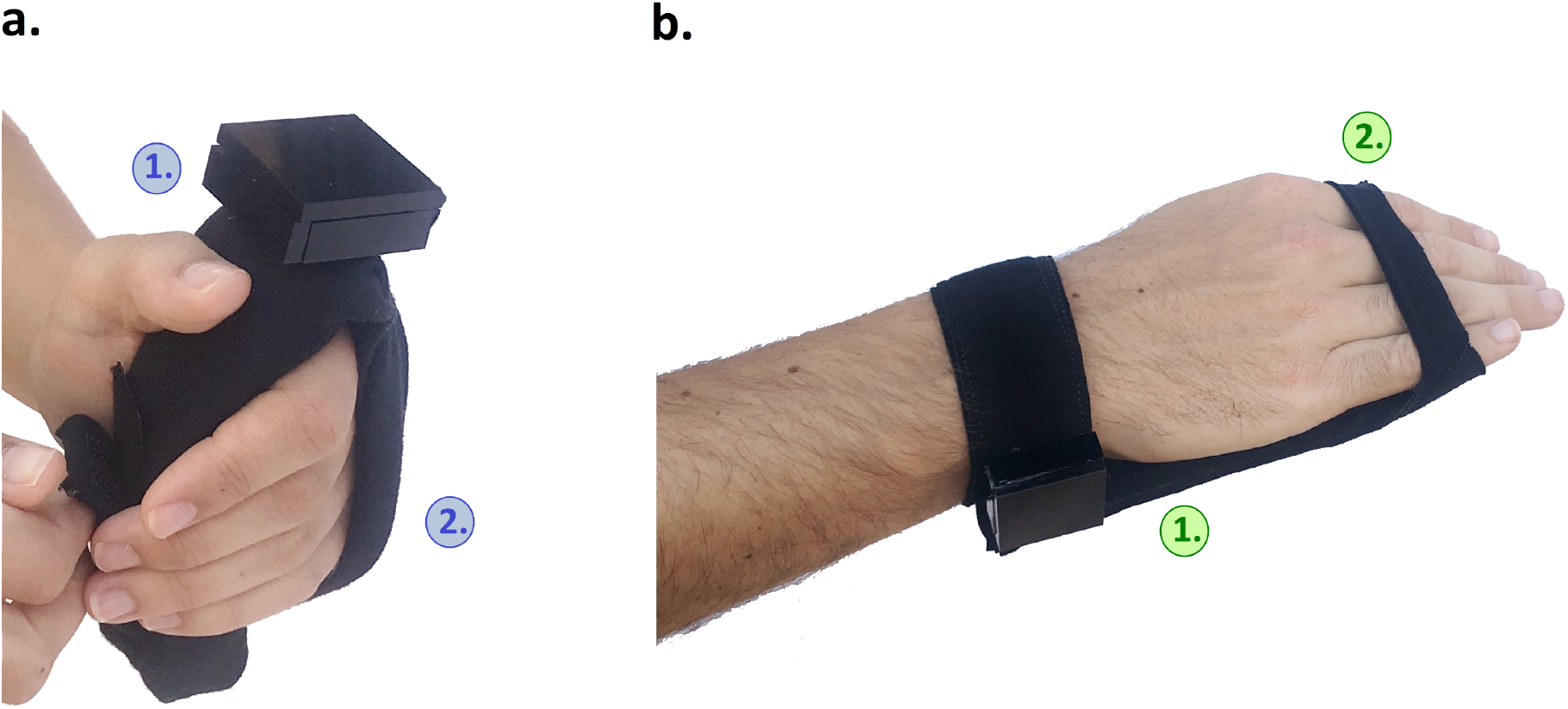
VTS Palm version V1 and version V4. **(a**.**) 1**. The affected hand can be placed around the rod-shaped device. **2**. An adjustable strap slips over the fingers. **(b**.**) 1**. The brace-like form factor contains foam which conforms to the palm when fingers are extended or flexed. **2**. A single strap secures the fingers.

### Procedure and Measures

Measures were chosen to provide data on each design goal. There were four types of measures used in this study: timed tasks, binary correctness measures, Likert scale questions, and interview transcripts. Interview guides were carefully designed according to qualitative study design principles [55, 56] to prompt insightful discussion without leading participants. Questions were standardized across prototypes and participants, and proctors could ask follow-up questions to gather more information. We used the SRQR reporting guidelines [57]. The procedures had the following structure:

- Don and doff the prototype (*baseline*)
- Don the prototype (*with experience*)
- ADLs while wearing the device
- Questions on physical and social comfort
- Doff the prototype (*with experience*)
- Likert scale survey

#### G1: Time to Don/Doff

Participants were first asked to don and doff the prototype. Their speed was measured using a stopwatch. They were not asked to rush. The proctor said “Could you please try putting on the device? Let me know when you are done by saying ‘I’m done.’ The proctor then asked the following questions:

*“What do you think?”*

*“Was it easy or difficult to get on?”*

*“How would you make it easier to don?”*

Next the proctor demonstrated how to don and doff the device using their own hand. The participant donned the device again and provided feedback on the challenges and successful features. Participants later doffed the prototype at the end of the session. These tasks were also timed using a stopwatch.

#### G2: Reliance on One Hand

After donning the prototype for the second time, participants were asked to demonstrate three tasks of daily life (writing on a clipboard, holding something in their lap, and walking) while wearing the prototype. These tasks and the don/doff tasks were used to reveal challenges and elicit participant feedback.

#### G3: Correct Use

Each time that participants donned the device, their correctness was recorded as a binary measure. To be considered correct, the device must be in the correct position and adjusted securely. After removing the prototype at the end of the session, participants were given a Likert scale survey to rate their level of confusion when using the prototype.

#### G4: Physical and Social Comfort

While wearing the device, participants were asked a series of interview questions about comfort that included both physical and social factors. Three examples are shown below.

*“What activities would you do while wearing this?”*

*“What do you think about wearing this in public?”*

*“Do any parts of the device cause discomfort?”*

The Likert scale questionnaire, given at the end of the session, also included three questions regarding physical and social comfort.

#### G5 and G6: Fit and Durability

Before doffing the prototype for the second time, participants were asked “How secure does the device feel?” The study proctor also recorded observations of fit, breakage, or strain throughout the study and these observations are reported in the discussion section.

### Data Analysis

Thematic content analysis was performed on the participants’ transcribed verbal responses. Between each round, transcripts of the interviews were read and coded independently by two of the investigators. These codes were then discussed and checked against the original transcripts. Next, actionable codes were discussed to determine revisions for next round. Revisions were made to address codes supporting a design change with no contraindications.

After three rounds of feedback, each investigator grouped codes to define themes of perspectives across all participants and prototypes. These themes were then discussed among the two investigators with reflection on potential bias from individual participants and/or the investigators. Don and doff times were averaged across participants for each round.

## Results

Participants were able to don all prototypes in less time than the VTS Glove: *VTS Phalanx* Mean = 57 s, SD = 19 s; *VTS Armband* Mean = 44 s, SD = 28 s; *VTS Palm* Mean = 49 s, SD = 26 s (with experience). Data from the timed tasks and correctness measures are provided in Figure 5. Feedback from each round and images of all prototypes (Versions V1-V4) can be found in Appendix A. All transcripts were analyzed and data are presented below.

**Figure 5.**
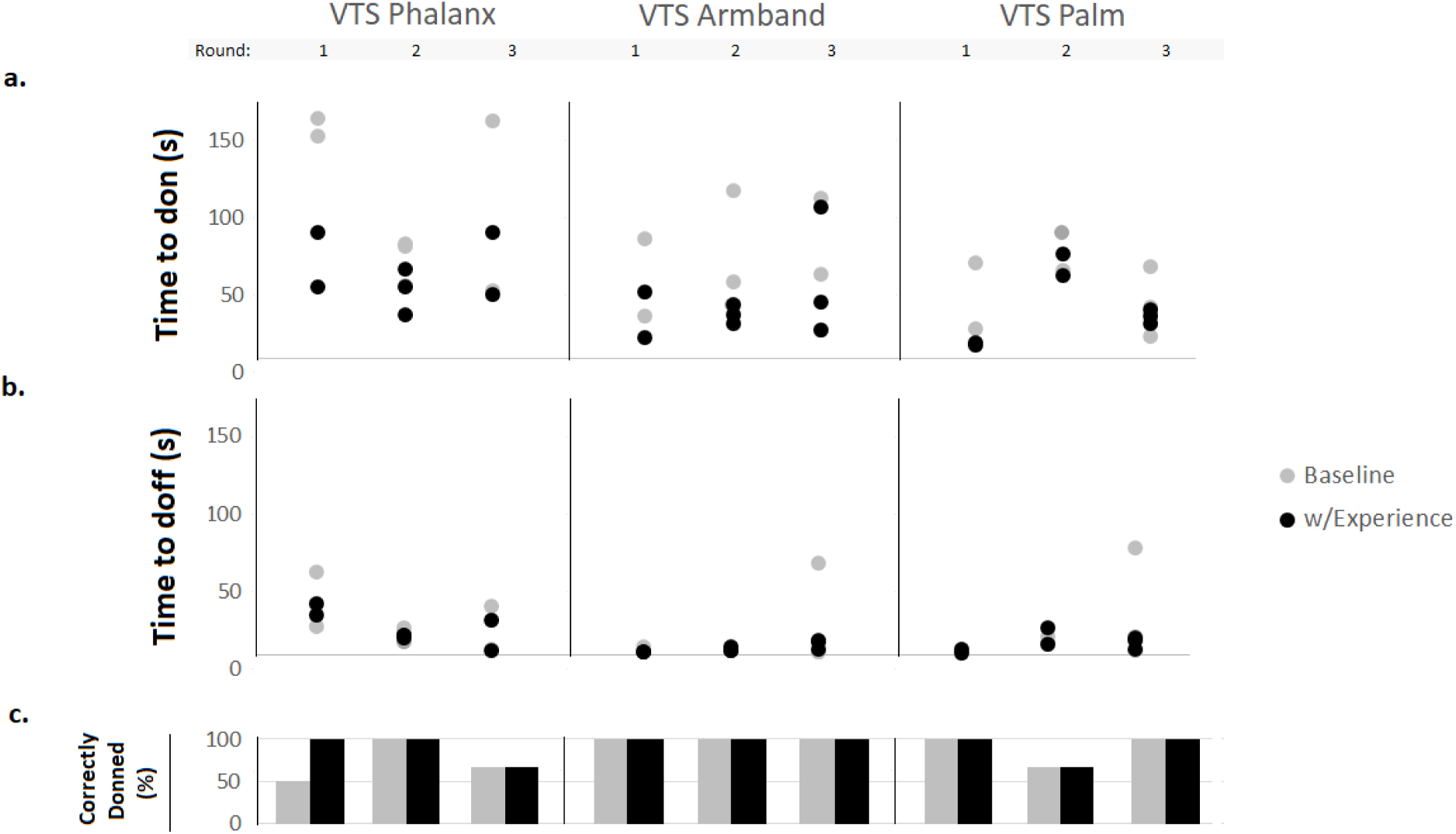
Quantitative data for each design in Study 2. **(a**.) Time required to don the prototype for each participant. Each set of vertically stacked circles represents one round in the study. **(b**.) Time to doff the prototype. **(c**.) Percent of participants who correctly donned the prototype for each round and each device.

### VTS Phalanx

The final VTS Phalanx (Figure 2b) uses a t-strap design to connect the fingers and distal forearm, so that more of the hand is uncovered. Electronics can be mounted securely at the wrist, and actuators can be mounted at the fingers. Two straps attach the design to the hand: one at the fingers and one at the wrist. This adjustable strap can be stretched over the fingers in one movement with the anterior arm facing away, not requiring supination or extension of the fingers. The strap at the wrist attaches using an adjustable side release buckle.

### VTS Armband

The VTS Armband (Figure 3b) was revised to be more narrow than its preliminary design, at 2 inches. At one end of the armband is a cinch buckle that allows the armband to be secured using one hand. The buckle frame is 0.75 inches wide. At the opposite end of the armband is a tapered point. Fabric is layered to make this tapered end rigid. This end secures using Velcro.

### VTS Palm

The final V4 version of the VTS Palm (Figure 4b) has a brace-like form factor that conforms to the palm using 1 inch foam embedded in between two fabric layers. This pad can be worn with the fingers extended and relaxed, or can be gripped by fingers that are flexed. Two straps attach the prototype to the hand. One strap at the wrist wraps around the arm and attaches using Velcro. A second strap stretches over the fingers.

#### Perspectives

#### Hand supination was challenging

Five out of eight participants expressed that they had difficulty flipping over their hand to reveal their palm (a motion known as supination). Proctors also noted that supination was impossible even for a participant with near normal function in their arm, P5.

These comments were made in response to questions about the donning process when evaluating three of the prototypes: one version of each design. Two of these prototypes included a strap that wraps around the limb, requiring the arm to rotate in pronation and supination (e.g. Figure 3a). The other prototype, version V2 of the VTS Phalanx design, included loops that stretch over each finger (Figure 6a). This fixture encouraged donning in the supinated position particularly when the hand is curled (due to spasticity), making the fingers only accessible with the anterior arm exposed. For example, participant 4 stated:

**Figure 6.**
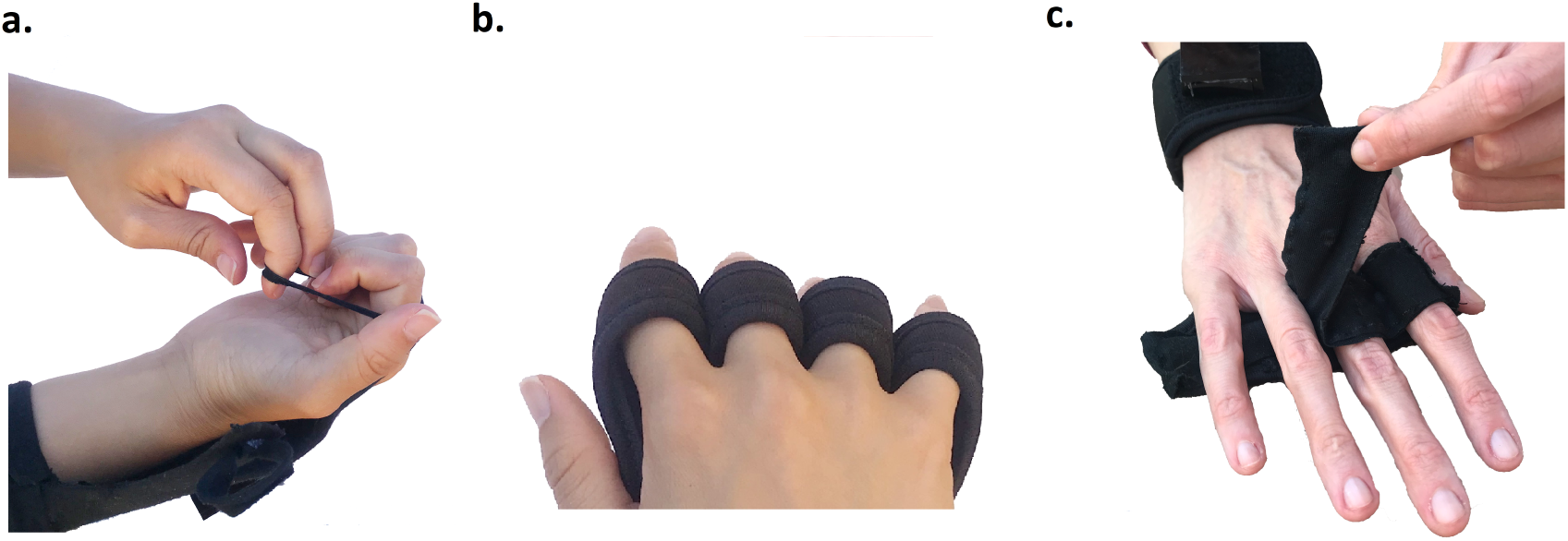
Design features that participants struggled with in Study 2. **(a**.) Participants struggled to supinate their arm and expose the volar hand to stretch loops over each finger. **(b**.) Participants could not insert their fingers into finger sleeves. **(c**.) When magnets were placed between the fingers, participants struggled with fit and finger abduction.

*“I can’t do the external rotation [supination]*.*” – P4*

Instead of wrapping a strap around the arm, later prototypes included a cinch buckle to allow participants to tighten straps in a cinching motion using just one hand.

#### Fingers may be flexed or open

VTS Palm V1 (Figure 4a) was designed to be gripped within the palm since participants with spasticity may have fingers in tight flexion. Commercial braces use this form factor, such as the Hand Contracture Carrot Orthosis (AliMed, Inc.). It is known that spasticity varies in different physiological conditions [58] and participants explained how this variation in spasticity impacts device design. Participants described that, due to changes in spasticity, their fingers may be either open or flexed without their awareness throughout the day. Thus, a rod-shaped form may not make reliable contact with the hand at all times. Subsequent prototypes of the VTS Palm design (V2-V4) used a flexible, brace-like form factor that makes reliable contact whether or not fingers are in flexion.

#### Fabric coverage may impact comfort

In answer to questions about comfort, participants also discussed sweat. The VTS Phalanx design and the VTS Palm design were associated with some discussion on fabric coverage and sweat, including how it relates to sanitation.

*“With the palm being one of the areas that is more sensitive to heat, sweat is a concern*.*” – P1*

#### Separate finger attachments prevent fit and require dexterity

Two participants were unable to don a prototype in the study. In both cases, the prototype (VTS Phalanx V3 and VTS Palm V2) included individual attachments for each finger. The remainder of participants who were able to fully don the prototype expressed difficulty with the interaction.

VTS Phalanx V3 used magnets between each finger that allowed the strap to “snap” on (Figure 6c). Large hands did not accommodate any fixtures between phalanges, and participants could not abduct their fingers to accommodate the magnets without help from the able hand. Participant 5 described this concisely: *“Spreading fingers is very difficult to do*.*” – P5*

VTS Palm V2 attached by resting the palm on the table and slipping each finger into a sleeve (Figure 6b). Though this design did not require the fingers to be held open (instead using a table or surface to extend the fingers), these sleeves required the fingers to be inserted. Participants, especially those with flaccid paralysis, struggled to slide their affected fingers or arm forward, which was necessary to insert the fingers to don this prototype. Participants who were able to fully don this prototype measured nearly double the donning time as the other versions of this device (Mean V2 = 76.5 s vs. Mean V1 = 44.7 s and Mean V3 = 41.0 s without experience). There was also a difference in donning time when donning with experience (Mean V2 = 67.0 s vs. Mean V1 = 9.5 s and Mean V3 = 20.0 s).

#### Social comfort is perceived as less important

63% (5/8) of the participants made comments to the proctors suggesting that they have diminished concern for public opinion. This view was often restated – when asked about their level of social comfort while wearing the prototype, or about how they would design the device. Some participant comments include:

*“I don’t care what people think when they see it. Maybe it’s me, I’m not a shy person.” – P1*

*“Ever since this stroke, I stopped caring what people think when they look*.*” – P5*

*“I’m not interested in what’s cool. Whatever works for my health, that’s cool*.*” – P8*

However, participants did make aesthetic requests. Namely, the request from four participants that the VTS Armband be more narrow so as to look more like a smartwatch. In addition, half the participants responded with lower ratings of social comfort when in public versus around friends and family (Likert scale ratings are graphed in Appendix A). Fisher’s Exact Test did not show a significant difference between these ratings across the prototypes (p = 0.59), but the skewness are different between the groups (Family: −1.7; Public: −0.85).

#### Affected hand is infrequently used

All but one participant (7/8) in the study remarked that they do not use their affected hand. Such statements included:

*“I can’t use this hand right now anyways.” – P2*

*“I don’t do anything with this arm.” – P8*

## Discussion

### Study 1

The VTS Glove successfully delivered wireless, wearable stimulation over eight weeks. Hardware dimensions were minimized to allow components to fit on the back of the hand, and the low-profile hardware was able to deliver over four hours of battery life, enable sensing, stimulation, and data collection. A limited selection of small vibration motors are available, and the glove’s hardware platform could be used to power a variety of these actuators, while the onboard program can be changed to apply a variety of stimulation patterns. Damage during the eight week period was limited to peripheral wires, which suggests that the central circuitry is robust.

Wearing the device for numerous hours each week appears feasible, based on feedback and participation time logged to the device. Participants with vibration enabled on their device each received over 140 hours of stimulation. Adherence under real-world conditions cannot be forecasted based on these data. However, these data provide supportive evidence that such a device is tolerable with or without vibration enabled. Participants mostly reported wearing the device while doing restful activities (59.6%), although they also reported wearing the VTS Glove to meals, movies, church and other gatherings. These data suggest that the glove was mobile and socially comfortable.

Some participants experienced significant difficulty when donning and doffing the VTS Glove. Such difficulty could result in strain and frustration, and may dissuade prospective users from adopting a new technology [35].

### Study 2

The primary vulnerability of the VTS Glove was its form factor, which was difficult to don in the presence of diminished hand function. Study 2 exposed ways to improve this interaction. Transcript data was used in conjunction with Likert scale ratings and quantitative measures to add triangulation to the findings.

#### G1: Time to Don/Doff

New VTS prototypes were designed to improve donning time by including accommodations. The prototypes did not require extension of insertion of the fingers, and all designs were found to require less time for donning/doffing than the original VTS Glove. It is known that contracted fingers and limbs can make dressing difficult due to their inability to stretch and flex; however, we noted that the limpness of flaccid fingers also makes don and doff challenging.

#### G2: Reliance on One Hand

Participants had to manipulate their affected hand using their unaffected hand. Supination of the affected hand was not achieved by some participants. To expose the palm, participants had to grip and twist their affected arm with their unaffected hand; this left their able hand fully occupied. As a potential solution, devices can attach at the dorsal hand or without view of the anterior arm. Participants can grip their affected fingers, stretch them open and rest the affected hand on the prototype (e.g. Figure 4b). Their able hand is then free to secure the device on the dorsal side.

Pronation may be similarly difficult for some patients. Survivors with weakness or spasticity may struggle to rotate their forearm in either orientation. Attachment using a cinch buckle allows patients to tighten devices securely, without pronation or supination, using one hand.

#### G3: Correct Use

Most designs were associated with high percentages of correct donning, and low ratings of confusion on Likert scales. However, novel features such as magnets were confusing to participants.

#### G4: Physical and Social Comfort

Sweat was a concern for participants. Therefore, later versions of the prototypes minimized fabric coverage to address this concern. Related to social comfort, some participants reported that they do not care about the opinion of others. However, the same participants made aesthetic requests and reported different ratings on their level of social comfort with friends and family versus the public. Despite reports that they do not care about the opinion of others, social comfort may be a concern.

#### G5: Fit

Certain design choices may enable improved fit. Designs that included individual finger sleeves or attachments between the fingers were not able to fit all participants. Buckles helped participants tighten the device. Final designs are ambidextrous.

#### G6: Durability

Proctors observed little strain of the prototypes, perhaps because these designs did not require insertion of the fingers or limb. The VTS Glove from Study 1 required repeated tugging and stretching that causes wear. Participants with limited dexterity have gross movements and therefore can cause device strain. More accessible designs may reduce interaction with the device and thus reduce damage.

## Conclusion

We designed the VTS Glove, a wearable device to provide vibrotactile stimulation to the affected hand in chronic stroke. In contrast to existing apparatus, the VTS Glove can provide stimulation for extended durations in a mobile form factor – to enable further study that expands on promising prior work regarding therapeutic vibrotactile stimulation. The glove is wireless, rechargeable, and includes small, dynamic hardware that can be programmed to provide stimulation at a variety of settings. Based on the result of an eight-week study, adherence to this method of therapy appears feasible. Participants each wore the device for over 140 hours, both at home and in social settings. The hardware was robust, but the form factor of the glove needed revision. Extension and insertion of the fingers was necessary to don the VTS Glove, and although these motions may be common in wearable device and garment interaction, they are a barrier for users with upper limb disability.

For clinical devices, efficacy depends on use. To reduce barriers to use and promote technology acceptance, we next performed a study to better understand factors influencing stroke survivors’ experience with an upper-limb wearable device. Results suggest that arm supination is another donning motion that is difficult for stroke survivors. Devices can be designed to avoid interaction in view of the anterior arm. Separate finger attachments, such as sleeves or loops, were found to require insertion of the fingers – once again challenging participants. Most survivors reported a reduced regard for social comfort, though ratings of comfort while wearing a device in public versus with family were differently skewed. Study 2 also provided the opportunity for revision of the new device designs (VTS Phalanx, VTS Armband, and VTS Palm) which all can be donned in an average time of 48 seconds (vs. 5.05 minutes to don the VTS Glove).

## Declarations

### Ethical Approval and Consent to participate

Study 1 was approved and overseen by the Office of Research Integrity’s IRB for the Georgia Institute of Technology. Study 2 was approved and overseen by the IRB of Stanford University. All participants provided written consent before beginning the study. ClinicalTrials.gov Identifier: NCT03814889.

### Consent for publication

Not Applicable.

### Availability of supporting data

Not Applicable.

### Authors’ contributions

AMO, TES, CES, MGL, and KF contributed to the study design and methodology. CES and BR designed interview guides and fabricated the prototypes used in this research. CES, MGL, and KF screened participants. CES and BR transcribed and analyzed data, then consulted with other authors. All authors drafted, edited, and approved the final manuscript.

## Acknowledgements

Not Applicable.

