## Appendix A for "Perspectives on the design and performance of upper-limb wearable stimulation devices for stroke survivors with hemiplegia and spasticity"

### RESEARCH

#### Appendix A

##### Introduction

This document contains details about the wearable device prototypes and participant feedback in each round of Study 2.

##### VTS Phalanx design

Versions of this design all attached at the distal forearm and the fingers. Electronics can be mounted at the wrist or on the back of the glove, while vibration motors should be embedded in contact with each dorsal proximal phalanx. Laser-cut and 3D printed components formed the basis for all versions of this design, which eliminated the need to use a commercial component as the base layer.

*Round 1*

This design shares the appearance of a fingerless, palmless glove. However, the fingers are attached using two flexible straps rather than individual sleeves (Figure A.1a). These straps enable donning without extending the fingers. Each dorsal proximal phalanx fits into a rigid arch where actuators or sensors can be placed. Arches are mounted on a foam bar inside the garment that flexes to accommodate both large and small hands. The palm was kept uncovered for cleanliness.

Participants described this design as comfortable and secure, as shown in the table of thematic codes in Figure A.4a. However, it was noted that the amount of fabric may lead to sweating.

*"It felt like part of my hand." - P1*

*"This is probably the easiest glove I have tried to put on" - P2*

*Round 2*

This version (Figure A.1b) reduced the amount of fabric from Round 1 by making the fabric narrow and hand-shaped. This prototype attached to the hand using a loop over each finger. Rather than two straps (used in Round 1), five straps aim to make the design more intuitive. The stretchable loops were designed to be tight on the finger, while being able to entirely stretch over a contracted finger without requiring extension.

This version, even more so than the prior version, required participants to supinate their forearm while donning. Proctors observed that supination was impossible even for participants with near normal function in their arm. Two of the three participants in this round verbalized difficulty with this motion (Figure A.4a).

*"I cannot do the external rotation" - P4*

*Round 3*

To eliminate the need for supination of the hand, the design was flipped to attach at the dorsal hand rather than volar hand. In theory, participants may stretch open their affected hand using their able hand, and place the affected hand to rest on the device. The weight of the affected hand then holds the device in place while the able hand secures the closures. To attach at the fingers, this design used magnets between each phalanx and allowed the strap to "snap" on. Figure A.1c shows the magnetic strap midway through attachment or removal.

Though attachment using magnets required almost no movement from the participants, large hands simply did not accommodate any fixtures between phalanges. Participants considered the device comfortable, but also expressed that the design was "confusing" and "difficult to don" (Figure A.4a). Using magnets as a fixture confused participants. On the Likert scale survey (Figure A.3a), participants using this design agreed that "This device was confusing to don." Participants did not report difficulties related to supination when interacting with this design.

*The magnet strap confuses me." - P4*

*"Spreading fingers is very difficult to do." - P5*

*Final Design*

In view of participant feedback, a final design was created. This version included the *least amount of fabric* to cover the hand – using a t-strap design to connect the fingers and distal forearm. To prevent confusion and *reduce steps in the donning process*, the design is attached to the fingers using a single strap. This adjustable strap can be stretched over the fingers in one movement, *not requiring supination or extension of the fingers.*

##### VTS Armband

This design was a band which wrapped around the forearm. Laser cut components form the basis of the

Full list of author information is available at the end of the article

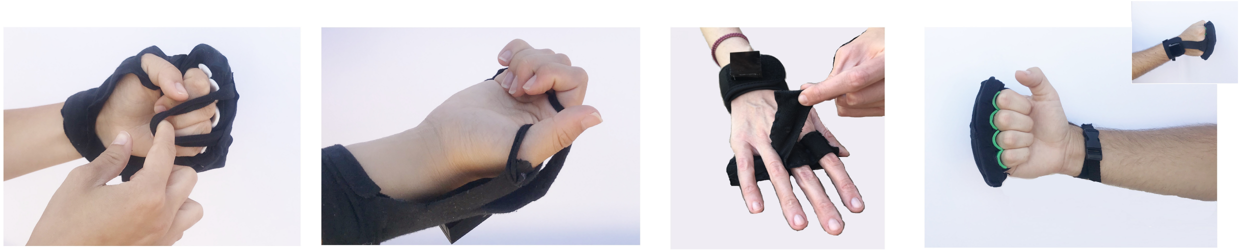

a. b. c. d.

**Figure A.1** Versions of the VTS Phalanx device through rounds 1-3 and a final version. During the design study, the amount of fabric bulk was reduced and attachments were simplified to allow donning without forearm supination.

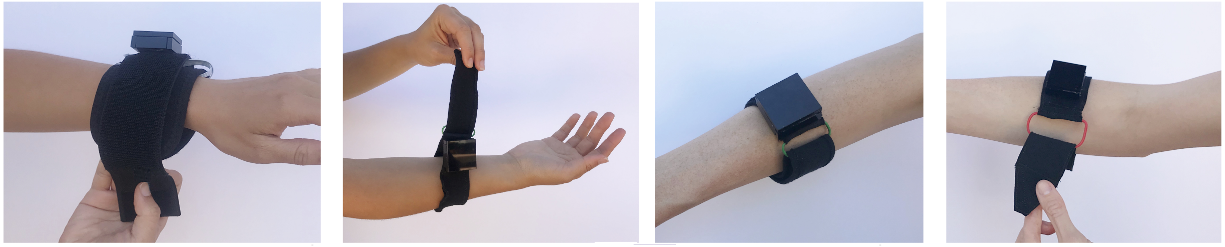

a. b. c. d.

**Figure A.2** Versions of the VTS Armband through rounds 1-3 and a final version. A cinch buckle was added and enlarged to enable one-handed tightening of the device.

prototypes. Electronics can be mounted along the band, where vibration motors should be embedded to apply stimulation to the forearm.

*Round 1*

This design consisted of a padded fabric band (3.0 in. wide) attached by a Velcro strap (total length: 15 in.). This length was chosen to accommodate arms of different sizes. Figure A.4b displays participants' feedback about the armband design. According to interview feedback, participants struggled to supinate their arm when attaching the Velcro of this initial armband design. The orientation of the hook-and-loop was also a concern of participants when the abrasive hook-side of the Velcro touched the skin. Proctors observed that the armband would swing, fall or hit nearby furniture when the Velcro was released. Both participants mentioned an aesthetic request – that the strap be more narrow so the device could look more like a smartwatch.

*"Make the band thinner. Then people might mistake it as a smartwatch." - P1*

*Round 2*

This version was designed to be more narrow than Round 1 (1.5 inches). Like the prior version, the band attached using Velcro, and a cinch buckle was added. The objective of this buckle was to allow participants to tighten the device in a cinching motion using just one hand. This would eliminate the need to supinate the forearm. The buckle also made removal of the VTS Armband more easy: reducing the requisite strength of the Velcro needed to hold the device closed, and preventing the Armband from falling off the arm when the Velcro is released. Velcro no longer comes in contact with the skin using this design. When this design was tested, participants were not observed trying to supinate their forearm; however, participants reported difficulty when trying to insert the strap into the buckle.

*"It was a little hard getting the end of the strap through the loop because it barely fits due to the width of the strap at the end." - P3*

*Round 3*

This version was widened slightly (2 inches), which provides better grip on the arm to prevent the band from slipping down. The buckle's frame was made more accessible by expanding its size to 0.75 inches. Figure A.2b-c shows the increased size of the buckle's frame. All participants reported comfort using this design, but one still reported difficulty inserting the strap into the buckle.

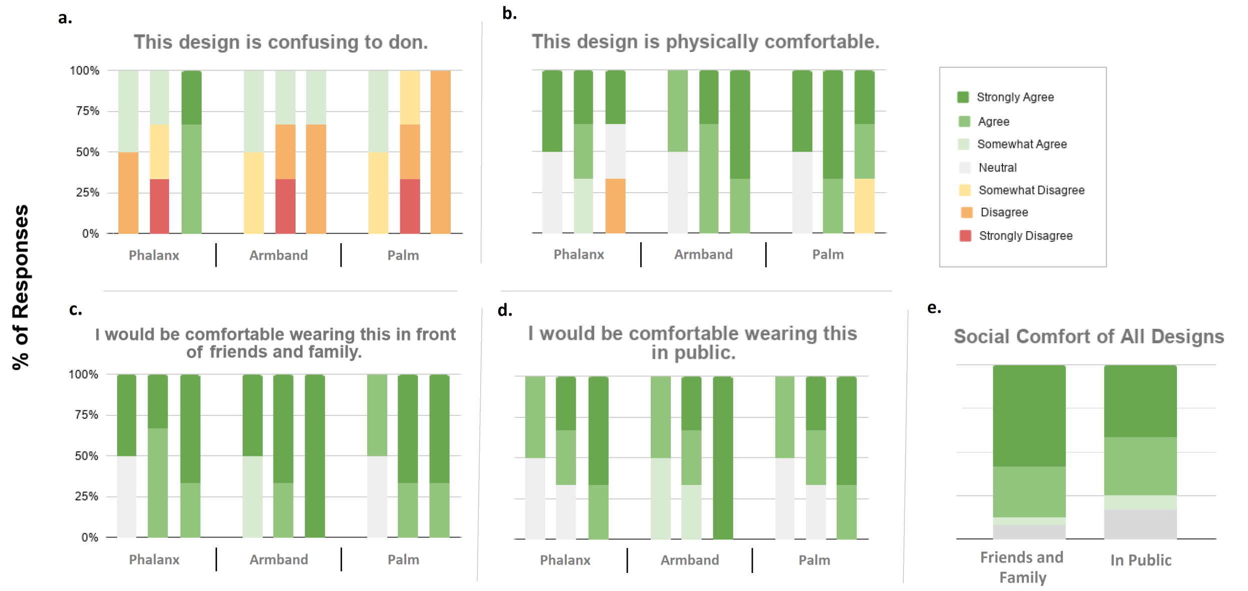


**Figure A.3** (a.-d.) Likert scale ratings for each device over three rounds. Each vertical bar represents one round. (e.) Total responses compared between statements about social comfort ("I would be comfortable wearing this [device] in front of...").

*"Getting the strap through the buckle was like threading a needle." - P6*

*Final Design*

A final design was made after the third round of participant feedback. This version (Figure A.2d) maintains the 2-inch-wide band and large buckle design, while adding one accommodation. The end of the band was tapered and ridged; forming a point that could be more easily inserted into the buckle using one hand.

#### VTS Palm

This design is intended to be grasped or attached to the inside of the hand. The hand may be in a flexed position due to spasticity, thus, the VTS Palm device is either cushioned or flexible. When finalized, electronics can be mounted at the wrist, and vibration motors embedded to make contact with the palm.

*Round 1*

This version is designed to be gripped within the palm. The device has two main parts: the padded rod and the elastic strap. The padded rod consists of a plastic tube embedded in a foam bar, to provide a combination of rigidity and comfort. The elastic strap secures the device to the hand. Participants can place their affected hand over the foam bar and adjust the strap.

During the study, participants agreed that this design would get in the way of activities (Figure A.4c). They also described that, due to changes in spasticity, their hand may be either open or flexed without their awareness throughout the day. Thus, this design may slip out of their grip.

*"It kind of puts this hand out of commission." - P1*

Participants also discussed sweat.

*"With the palm being one of the areas that is more sensitive to heat, sweat is a concern." - P1*

*Round 2*

To prevent the prototype from slipping off the hand, this version used a semi-flexible brace-like form factor that is compatible with both flaccid and flexed hands. This design did not include significant bracing structure, but one could reinforce the design to splint the hand. The donning process is very similar to the prior design: the participant opened the fingers of the affected hand with their able hand, placed their affected hand over the prototype and secured the attachments. The prototype was attached at the distal forearm with Velcro. The fingers slip into loops attached to the top of the prototype.

During the study, participants struggled to insert their fingers in the loops. These loops did not fit the fingers of some; participant P3 was unable to don this device and provide feedback on its comfort. Participants with flaccid paralysis struggled to slide their affected fingers or arm forward, which was necessary to don this prototype.

| | a. VTS Phalanx | | | b. VTS Armband | | | c. VTS Palm | | |
|---|---|---|---|---|---|---|---|---|---|
| | 1 | 2 | 3 | 1 | 2 | 3 | 1 | 2 | 3 |
| Device is easy to don | 50% | 100% | | 100% | 67% | 67% | 100% | | 100% |
| Device is secure | 100% | 33% | | 100% | 67% | 67% | 50% | 33% | 33% |
| Device is lightweight | | 33% | 33% | 50% | 67% | 33% | 50% | 33% | |
| Device is comfortable | 100% | 33% | 33% | 100% | 100% | 100% | 100% | 67% | 33% |
| Device would not limit activities | 100% | 67% | 67% | 100% | 100% | 67% | | 33% | 33% |
| Comfortable in public | 50% | 67% | 100% | 50% | 100% | 100% | 50% | 33% | 100% |
| Device is difficult to don | 50% | | 67% | | 33% | 33% | | 33% | |
| Device is confusing to don | 50% | | 100% | 50% | 33% | 33% | 50% | | |
| Device is too bulky | | 67% | | | 33% | | | 33% | |
| Device may cause sweating | 50% | 33% | 33% | | | | 50% | 33% | |
| Device would limit activities | | | | | | | 100% | 33% | 33% |
| Uncomfortable in public | | 33% | | | | | | 33% | |
| Difficult to supinate arm | | 67% | | 100% | | | | | 33% |
| Design change: change strap colors for visual clarity | | | | | | | 50% | | 33% |
| Design change: reduce band thickness | | | | 100% | 33% | 33% | | | |
| Not concerned with public opinion | 50% | 33% | 100% | 50% | 33% | 100% | 50% | 33% | 100% |
| Affected hand is rarely used | | 33% | 67% | | | 33% | 50% | 67% | |

**Figure A.4** Interview codes for each design across all three rounds. A colored block indicates the fraction of participants who made a response including that code. Codes found two or more times throughout the study are included. Codes are grouped into three themes for presentation clarity: positive codes (green), negative codes (red), and codes about lifestyle (blue).

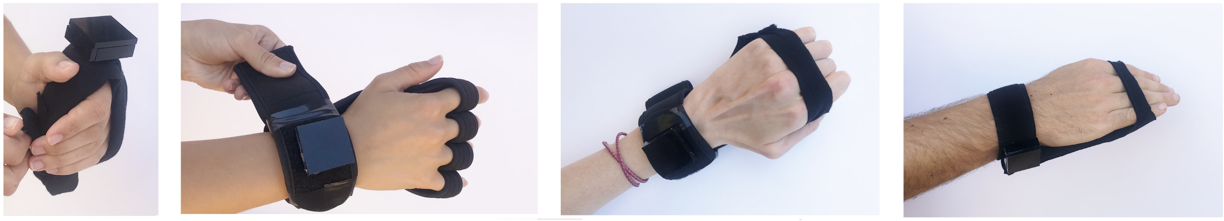

a. b. c. d.

**Figure A.5** Versions of the VTS Palm device through rounds 1-3 and a final version. A graspable rod form factor was changed to a brace-like form factor that is compatible with clasped or relaxed hands.

*Round 3 and Final Design*

This design shared the brace-like form factor of the prior version, but replaced the individual finger attachments with one flexible strap. This strap aimed to eliminate challenges with fit on different sized fingers, and allow participants to stretch over all the fingers without much manipulation. Feedback on this design was more positive, with all participants in this round reporting that it was easy to don with or without experience (Figure A.4c). One participant discussed that the device could feel more securely attached, and this was reflected in their ratings of physical comfort on the Likert scale survey (Figure A.3b). The final design made few changes from this version.
